# Rate of immune complex cycling in follicular dendritic cell determines the extent of protecting antigen integrity and availability to germinal center B cells^1^

**DOI:** 10.1101/2020.11.27.401315

**Authors:** Theinmozhi Arulraj, Sebastian C. Binder, Michael Meyer-Hermann

## Abstract

Follicular Dendritic Cells (FDCs) retain immune complexes (ICs) for prolonged time periods and are important for germinal center (GC) reactions. ICs undergo periodic cycling in FDCs, a mechanism supporting an extended half-life of antigen. Based on experimental data we estimated that the average residence time of Phycoerythrin-ICs (PE-ICs) on FDC surface and interior were 21 and 36 minutes, respectively. GC simulations show that antigen cycling might impact GC dynamics due to redistribution of antigen on the FDC surface and by protecting antigen from degradation. Antigen protection and influence on GC dynamics varied with antigen cycling time and total antigen concentration. Simulations predict that blocking antigen cycling terminates the GC reaction and decreases plasma cell production. Considering that cycling of antigen could be a target for the modulation of GC reactions, our findings highlight the importance of understanding the mechanism and regulation of IC cycling in FDCs.

## Introduction

Follicular dendritic cells (FDCs) form a dense network of cytoplasmic extensions in the B cell follicle (1–3) and are known for their ability to retain native antigen for a long period of time (1, 4, 5). FDCs were found to originate from perivascular mural cells in the spleen (6) and marginal reticular cells in the lymph node (7). Antigen is distributed mainly in the form of immune complexes (ICs) non-uniformly on FDC processes (5). FDCs bind antigen by CR1/2 (8) and/or FcγRIIB receptors (9), depending on activation and availability of complement proteins (10). While antigen is rapidly cleared from different body sites following immunization (11), long term antigen retention capacity of FDCs was believed to be due to a mechanism which protects antigen from damage (5). However, it was not understood how the ICs retained on FDC surface was maintained intact and stable for a long period (5).

Phan et al., discovered that ICs captured by subcapsular sinus macrophages, are transferred to follicular B cells and in turn FDCs acquire these ICs through complement receptors (12). Heesters et al., found that ICs acquired by murine FDCs are rapidly internalized and they reappear on the FDC surface (13). Experiments showed that ICs undergo multiple rounds of cycling in FDCs with significant amount of antigen remaining undegraded (13). Further, cycling of ICs was blocked by cytochalasin, an actin inhibitor, but the exact mechanism of IC cycling in FDCs is unknown (13). This finding suggested that cycling of ICs could be a process explaining the long-term antigen retention ability of FDCs. Cycling in FDCs was also found to be involved in protecting infectious agents such as HIV virions and FDCs could act as a constant source of HIV virions for infecting Tfh cells (14). While the timescale of IC cycling is unknown, it has been shown with nanoparticle tagged antigen, that particle size could determine localization of antigen in FDCs (15). Larger size particles preferentially localized on FDC surface and comparatively smaller particles are localized both in the interior and on the FDC surface (15).

FDCs are thought to be important for sustaining GC reactions and memory responses (16–19). GC B cells acquire antigen from FDCs in the form of ICs and this is believed to drive affinity maturation of B cells in the GCs (20). Importance of antigen availability in enhancing GC reactions has been demonstrated by several studies (21–23). Several accessory activities of FDCs other than IC trapping such as supply of BAFF (24–26) and maintaining follicular structure (27) and their importance are being studied (17). In addition to primary and secondary lymphoid organs, FDCs are also found in TLOs (Tertiary lymphoid organs) (28–30). Targeting FDCs in TLOs is considered as a promising strategy to disrupt such ectopic GCs (31, 32).

Mathematical modeling remains instrumental in explaining several biological processes. Particularly, *in silico* studies on endocytosis and recycling of various receptor molecules have provided insights into the mechanism, implications or timescale of the process and aided the interpretation of experimental data (33–37). In this study, we performed *in-silico* experiments and estimated the cycling timescale of ICs in FDCs for the particular antigen Phycoerythrin (PE). Using an agent-based model of GC reaction, we studied the implications of antigen cycling on GC reactions under different conditions.

## Methods

### Model for antigen cycling and estimation of cycling times

An agent-based approach was used to estimate cycling times where individual antigen particles undergo transition between two states – INTERIOR and SURFACE, corresponding to transition of antigen between the FDC interior and surface, respectively (Figure 1A). Transition times *t*_surface_ and *t*_interior_ were sampled independently from gaussian distributions *N*(*μ*_s_, *σ*_s_) and *N*(*μ*_i_, *σ*_i_) respectively.

**Figure 1:**
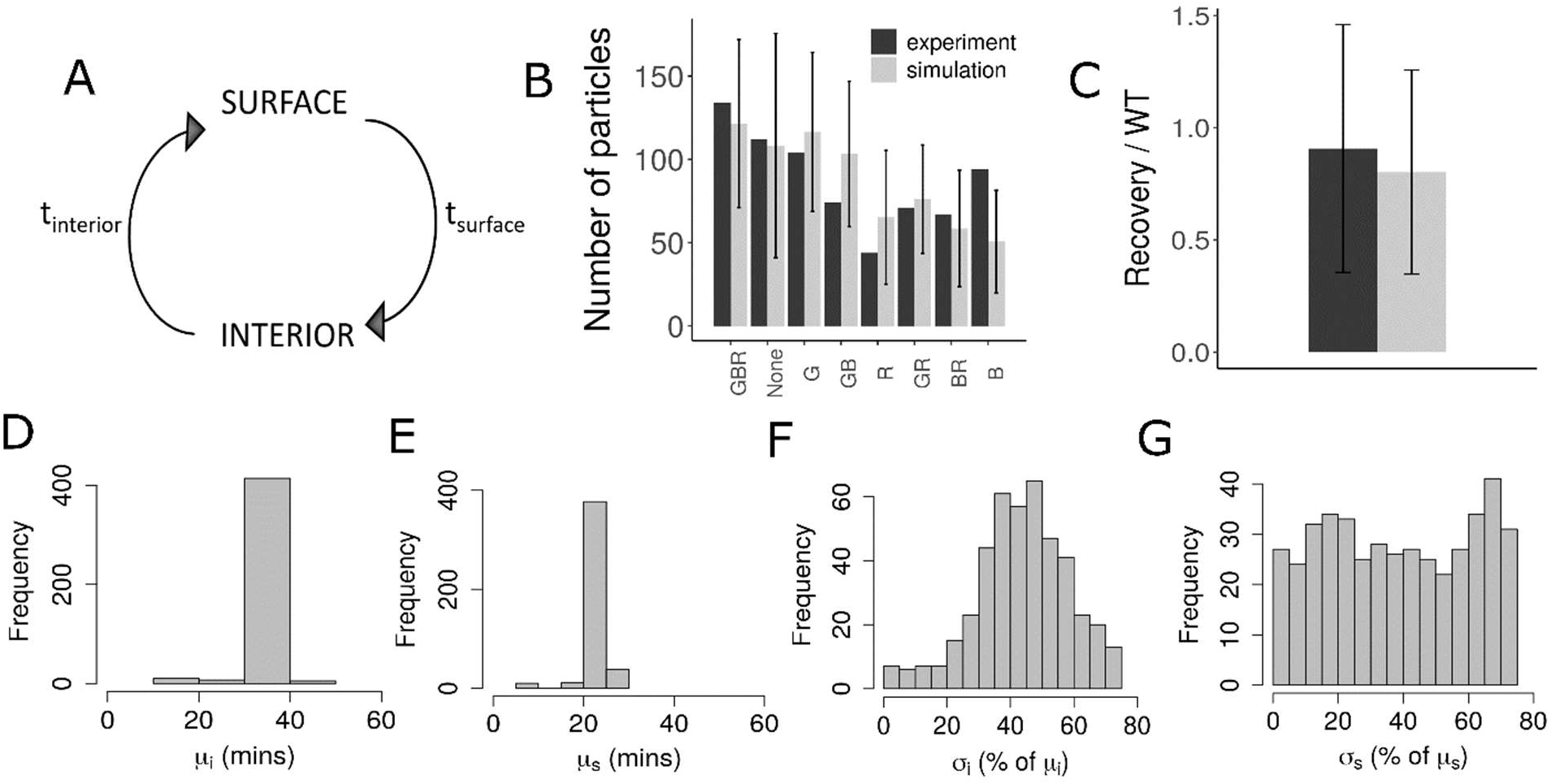
Estimation of PE-IC cycling time: A) Schematic representation of the model used to estimate cycling times. B) and C) Simulation of sequential staining and acid wash (explained in Methods section) compared to experimental data (13), for the parameter set with lowest cost, i.e. 21 and 36 minutes average time on surface and interior, respectively. B) Sequential staining of ICs with antibodies labelled Green (G), Blue (B) and Red (R) and the readouts show the number of IC particles labelled with the corresponding color combinations indicated below the bars. C) Acid-stripping of FDCs to remove ICs from surface followed by a recovery period. Readout represents the surface particle count after acid stripping and recovery, normalized with the surface particle count of control without acid treatment. The measured MFI of the recovery sample was normalized with MFI of the WT sample. D) – G) Parameter sets with Δ AIC < 2 with respect to the lowest cost parameter set. D) Average time in interior *μ*_i_, E) average time on FDC surface *μ*_s_, F) width of distribution for time in interior *σ*_i_ (as % of mean *μ*_i_), and G) width of distribution for time on FDC surface *σ*_s_ (as % of mean *μ*_s_). PE-IC: Phycoerythrin-immune-complex; *t*_surface_: time spent by ICs in state SURFACE; *t*_interior_: time spent by ICs in state INTERIOR; WT: Wild Type; MFI: Mean Fluorescence Intensity; AIC: Akaike Information Criterion; FDC: Follicular Dendritic Cell.

*In-silico* experiments (described below) were performed following the experimental setup in (13) and parameters *μ*_s_, *σ*_s_, *μ*_i_ and *σ*_i_ were varied to identify parameter sets with minimum cost with respect to the experimental results by an exhaustive parameter search. Parameters *μ*_s_ and *μ*_i_ were systematically varied between few minutes to 1 hour and *σ*_s_ and *σ*_i_ were varied between 0 – 80 % of the corresponding mean (*μ*_s_ and *μ*_i_, respectively). Cost of simulation results (*S*_k_) with respect to experimental results (*E*_k_) was calculated as follows:

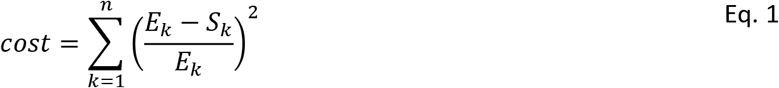

where *n* is the cumulative number of data points from all considered experiments. To set a cutoff for parameter estimates, parameter sets with Δ AIC < 2 with respect to the lowest cost parameter set were chosen for further consideration (Figure 1D-G).

#### *In silico* experiments

##### Sequential staining simulation

The simulation corresponds to the sequential staining of PE-ICs on FDCs in (13). In the experiment, the authors incubated FDCs with three distinctly labelled antibodies (with colors: green, blue and red) one at a time sequentially, to stain surface ICs. Each staining was performed with a particular labelled antibody for 5 mins, followed by a 60 mins period where no antibody is added. This resulted in ICs acquiring different combination of colors, depending on the localization of ICs during the time period of staining. Color combinations of PE particles were determined by visualizing the fluorescence of PE particles (13).

Addition of the labelled antibodies was simulated *in-silico* following the experimental setup (13) and antigen particles in state SURFACE were assumed to acquire the corresponding color of the antibody. The binding probability of staining antibodies was assumed to be 1, provided the antigen is in the state SURFACE at any time during the 5 minutes staining periods. As the color combinations of 700 PE-particles were examined in (13), 700 antigen particles were simulated and classified based on the color combination acquired as GBR, None, G, GB, R, GR, BR and B (G-Green, B-Blue and R-Red).

#### Acid wash simulation

The simulation follows the acid-stripping experiment performed in (13) to demonstrate the appearance of ICs on surface from interior of FDCs. The authors treated FDCs briefly with acid buffer which removes the ICs from the FDC surface and surface antigen was detected by quantifying MFI (Mean Fluorescence Intensity) of PE-staining, after a 30 minutes recovery period (13).

In the simulations, acid wash was performed by removing the antigen particles in the state SURFACE and then the system was allowed to recover for 30 minutes before determining surface particle count. Surface particle count determined after acid wash and recovery was normalized using the surface particle count determined in the absence of acid wash. Correspondingly, MFI data (13) of experiment was normalized with MFI from control (WT), to compare with the normalized surface particle count from the simulations.

As the initial distribution of antigen particles on FDC surface and in the interior at the start of the experiment is unknown, in both the simulations described above, particles were distributed randomly in the states INTERIOR and SURFACE using a probability sampled from uniform distribution. Simulations corresponding to each experiment were repeated 100 times.

### Germinal center simulations

We generated a new agent-based model of the GC reaction, which uses elements from previous state-of-the-art models (38–40) and was particularly developed to cover the dynamics of antigen on FDCs. This model includes a 3D discretized lattice, which is divided equally into a dark zone (DZ) and a light zone (LZ). For the affinity representation of B cells, a 4D shape space (41) is used, where the position of the B cell with respect to a predefined optimal position provides a measure for the antigen binding probability. Initially, 250 Tfh cells are randomly incorporated in the lattice. Tfh migrate and tend to accumulate in the LZ due to chemotaxis.

200 FDCs are distributed in the LZ. Each FDC has 6 dendrites that are each 40 micrometers long. Each FDC occupies more than 1 lattice site, hence there are several connected fragments distributed on several nodes for a single FDC. Rate constants of internalization and externalization for antigen are calculated from the average time spent on surface and interior of FDCs estimated from (13). For the simulations, average times of 36 and 21 minutes in the interior and surface were used and each FDC is loaded with 3000 or 1000 antigen portions. The antigen portions are distributed equally in different fragments of every FDC. Surface antigen amount on each FDC (*A*_surface_) was calculated by the following equation, where *A*_Total_ is the total antigen amount of an FDC, *k*_ext_ and *k*_int_ are the externalization and internalization rate constants respectively.

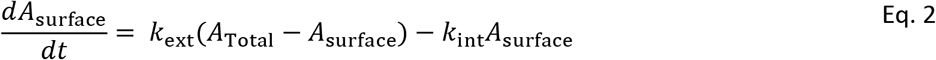

The calculated surface antigen amount is re-distributed on all fragments of the FDC. In simulations without antigen cycling, we used the surface antigen amount *A*_surface_ = *A*_Total_.

Degradation of antigen was modelled by a decrease in the surface antigen amount on FDCs over time as follows:

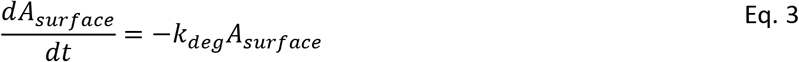

where 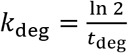 and *t*_deg_ is the antigen half-life.

Founder B cells with randomly chosen position and affinity are incorporated at a rate of 2 cells/hour for 96 h. Founder cells are assumed to divide 6 times before differentiating to LZ phenotype. During every cell division, B cells mutate with a probability 0.5 (42, 43) which would result in a shift in the shape space position.

LZ B cells collect antigen for a duration of 0.7 hours by establishing contact with the FDCs, depending on the binding probability of the B cell. Acquisition of antigen by B cell reduces the amount of antigen at a particular fragment of FDC. Failure to collect antigen within the fixed duration leads to a switch to the apoptotic state while successful antigen collection leads to the FDC selected state. FDC selected B cells establish contact with a neighboring Tfh cell and remains bound for 36 minutes. During this period Tfh signals only the B cell with highest amount of antigen collected. At the end of interaction time with Tfh, if the B cell has received signals for less than 30 minutes, it becomes apoptotic. B cells that received sufficient signals are selected, retain the LZ phenotype for 6 h and then recycle back to the DZ. Number of divisions of the recycled cell is determined by the amount of antigen collected with the maximum number of divisions limited to 6. After selection, mutation probability is reduced in an affinity dependent manner. 72% of the divisions are assumed to be asymmetric after which one daughter cell retains all the antigen. Daughter cells retaining antigen differentiate to output/plasma cells that exit the GC, while the other daughter cells acquire a LZ phenotype and proceed for the next round of selection. Steady state distributions of chemokines CXCL12 and CXCL13 are calculated and used for the chemotaxis of motile cells. Detailed assumptions of GC B cell dynamics and parameter values follow the supplementary text of (44) with the newly introduced dynamics of antigen in FDCs described here.

Simulations were performed using C++. ggplot2, plot3D and Bolstad2 packages of R were used for analysis and visualization of the results.

## Results

### Estimation of cycling time of PE-ICs in murine FDC

Periodic cycling of antigen in FDCs was demonstrated by sequential staining of PE-ICs on cultured FDCs with three distinctly labelled antibodies and quantification of the number of antigen particles tagged with different antibodies (13). The authors showed that upon acid stripping of ICs on FDCs and subsequent recovery, ICs reappear on FDC surface (13). The timescale of antigen cycling in FDCs is unknown as it is not directly measured in the experiments. However, as these experiments are informative in providing an estimate of the underlying timescale, we performed *in-silico* simulations following the same *ex-vivo* experimental protocol (see methods section). We used a model where antigen particles undergo transition between two states – INTERIOR and SURFACE (Figure 1A, detailed description in methods) and transition times (*t*_surface_ and *t*_interior_) follow gaussian distributions. We fitted the mean (*μ*_s_ and *μ*_i_) and width (*σ*_s_ and *σ*_i_) of the gaussian distributions of transition times to estimate the average time, IC clusters spend on the FDC surface and in the interior. We performed an exhaustive parameter search to identify the set of parameters with minimum combined cost to the experiments simulated. Parameter set with lowest cost indicated that particles spend an average time of 21 and 36 minutes on FDC surface and interior, respectively. The estimates were confirmed with a differential evolution parameter search over a larger range of parameters. Estimated parameter ranges are shown in Figure 1D-G. The width of the distributions estimated (*σ*_s_ and *σ*_i_) were broad (Figure 1 F and G). The experimental data could not be fitted with fixed cycling times (*σ*_s_ = 0 and *σ*_i_ = 0) suggesting that the variability in antigen presentation kinetics is critical to reproduce the data. In order to see whether the fit (Figure 1B and C) could be improved further, we tested various possibilities such as varying the initial distribution of particles in the two states, binding probability of staining antibodies and considering a bimodal distribution for the average time in the state INTERIOR. All these trials resulted in similar or higher cost values and could not improve the fit to the data. However, the deviation from the experimental mean for the number of particles labelled blue (represented as B in Figure 1B) could be due to large fluctuations in the behavior of antigen particles.

### Antigen redistribution on FDCs enhances GC reaction

To investigate if the cycling of antigen can influence the GC dynamics, we simulated the GC reaction (see Methods) in the presence and absence of antigen cycling (Figure 2). The mean transition times obtained as described in previous section were used to calculate the externalization and internalization rate constants *k*_ext_ and *k*_int_ in Equation 2. Due to cycling, antigen in FDC was distributed on the surface and the interior, with the distribution dependent on *k*_ext_ and *k*_int_. In the absence of cycling, all the antigen portions were distributed on the FDC surface. As a result, surface antigen amount on FDC was lower in the GC simulation with cycling when compared to simulation without cycling (Figure 2B). In the presence of antigen cycling, the GC volume at the peak of the reaction and thereafter (Figure 2A), was increased despite having lower surface antigen concentration. There were no marked changes in the affinity of plasma cells produced (Figure 2E). The changes in GC volume were reflected in the plasma cell production, showing an increased plasma cell production with antigen cycling (Figure 2D). Importantly, the lower surface antigen concentration did not impair GC reactions as antigen is subsequently replaced on the FDC surface at a timescale much shorter than the lifetime of GC reaction.

**Figure 2:**
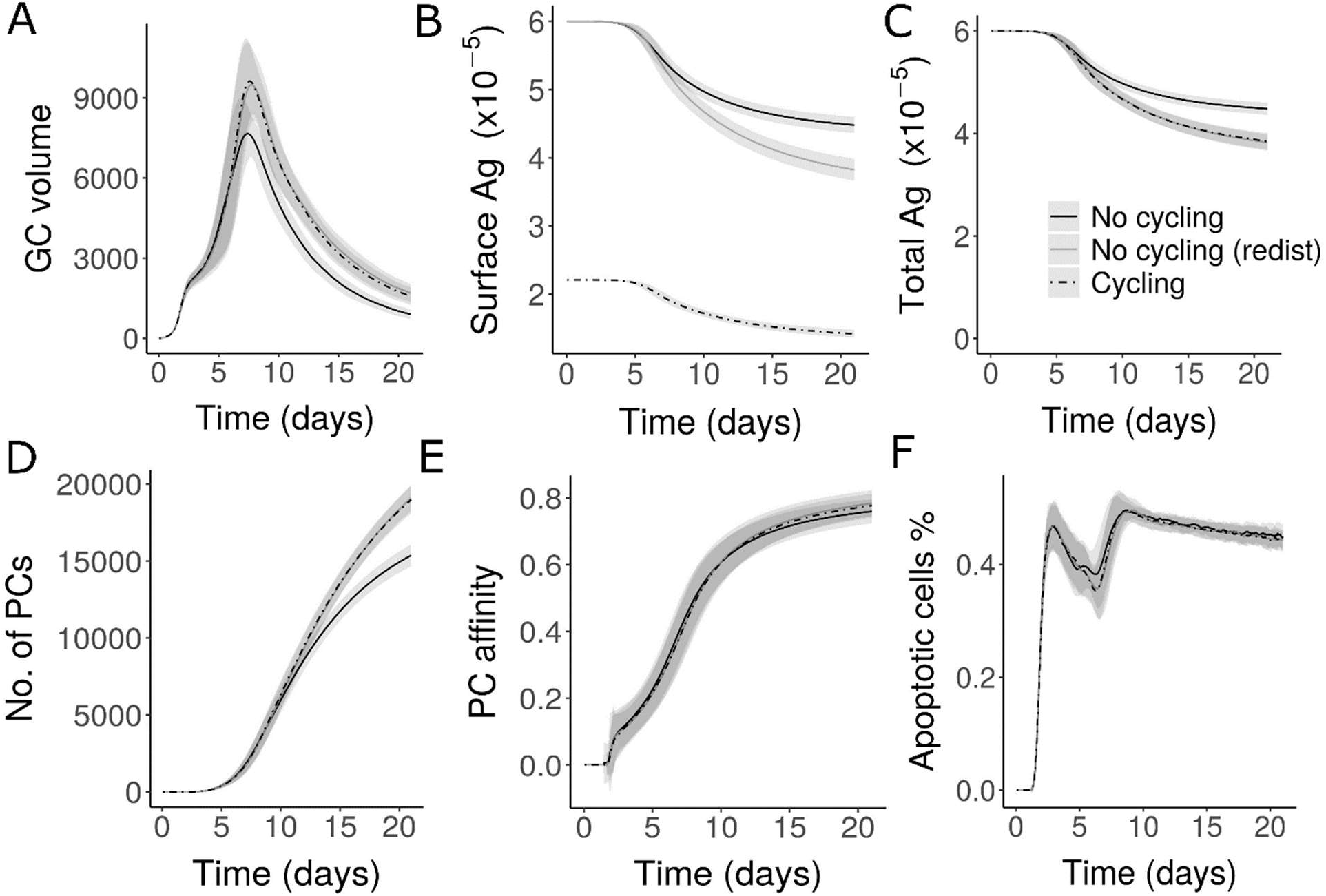
GC simulations in the presence and absence of antigen cycling: A) GC volume measured by number of GC B cells B) Surface antigen amount of all FDCs in the GC C) Total antigen amount including surface antigen and internalized antigen D) Number of Plasma cells produced E) Plasma cell affinity and F) Percentage of apoptotic cells. ‘No cycling (redist)’ represents simulations without antigen cycling (similar to ‘No cycling’) but includes surface re-distribution of antigen. Antigen amount is represented as number of antigen portions. GC: Germinal Center; FDC: Follicular Dendritic Cell.

In order to check if the enhancement in GC volume at later stages is due to the re-distribution of antigen from time to time, we re-distributed surface antigen at every time step in the simulations without antigen cycling (Simulations labelled ‘No cycling (redist)’). There were no differences in GC volume, apoptotic cells, number or affinity of plasma cells with cycling and without cycling in the presence of re-distribution (Figure 2), suggesting that the enhancement in GC volume and PC production seen are due to the re-distribution of antigen on FDC surface rather than a direct effect of antigen cycling. Hence, compared to the static representation of all antigen portions on FDC surface, re-distribution of antigen on FDC surface which can be brought about by antigen cycling can enhance the GC reaction.

### GC dynamics remains unaltered with varying cycling times

As there is a possibility that the cycling rates might vary dramatically depending on the particle size, we also varied the cycling times in order to check the impact on GC reactions. We chose the following average cycling times in the interior and surface of FDCs: (a) 10 and 50 mins – particles spend longer on surface (b) 36 and 21 mins – estimated from PE-IC data, slightly longer in the interior than surface and (c) 50 and 10 mins – particles spend much longer time in the interior of FDC. With varying cycling times, surface antigen amount is altered (Figure S1A). However, the GC dynamics, and output of GCs remained unaffected (Figures S1B-D). Similarly, when cycling parameters were varied in the estimated range, no major changes were seen in the kinetics of GC and output quantity or quality despite large changes in surface antigen amount (Figures S1 E-H). Small differences were observed between the different cycling times, in the simulations with lower antigen concentrations (not shown). This shows that GC reactions are robust against changes in cycling times provided the antigen concentration is not limiting.

### Cycling could impact GC reaction by limiting antigen degradation

Since the discovery of FDCs as long-term depots of antigen, it was not understood how easily degradable antigen is retained on FDC surface for a long period of time (5). It has been shown that constitutive endocytosis of vascular endothelial growth factor (VEGF) receptor protects the integrity of the receptor and blocking this process results in the shedding of VEGF receptors (45). Similarly, the antigen could be protected by cycling as this minimizes surface exposure. However, it is not clear how the antigen is protected from damage despite remaining available for GC reactions, hence we tested if antigen cycling could protect antigen efficiently by minimizing antigen exposure on FDC surface without impairing GC reactions. Antigen degradation was modelled using Equation 3 (methods section) assuming that the surface antigen amount is degraded with a fixed half-life. The amount of degraded antigen was dependent on the cycling times (Figure 3A) and the GC shutdown was delayed when the average time of antigen in the interior was higher (Figure 3C). Consequently, there is an increase in plasma cell production as well (Figure 3D). Thus, in a system with antigen degradation, antigen cycling can increase the GC reaction duration and limit antigen degradation, by this extending the time period of antigen availability for B cell selection. Longer time for antigen inside the FDCs was advantageous for the GC output in this case (Figures 3G-I). However, with low antigen concentration (Figures 3J-L), a longer residence time inside FDCs would dramatically decrease the amount of antigen available for GC B cells and hence, there is a trade-off between antigen protection and availability to GC B cells (Figure 3).

**Figure 3:**
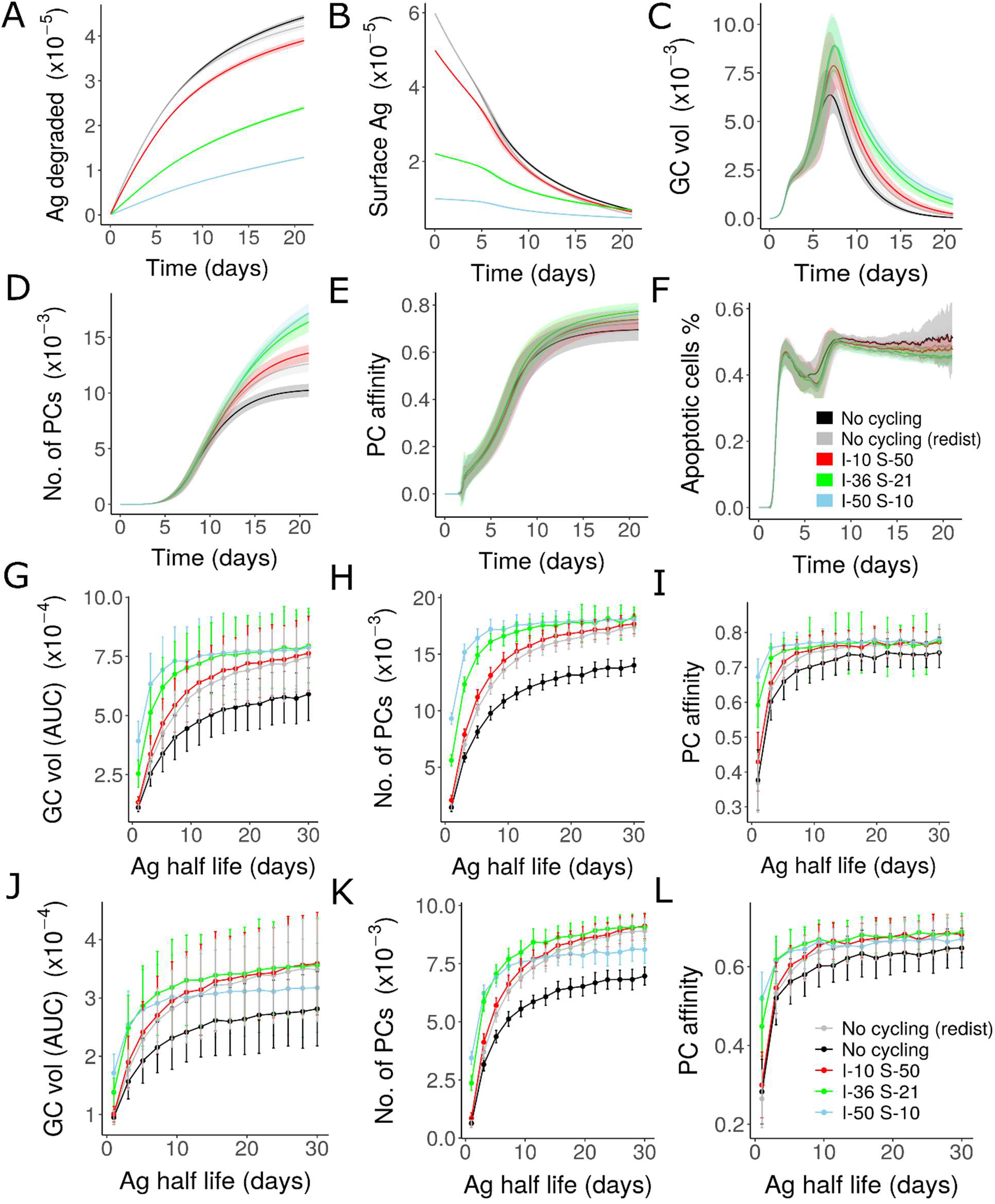
GC simulations in the presence of antigen degradation (Equation 3) with a half-life of 8 days: A) Amount of antigen degraded, B) Surface antigen amount, C) GC volume, D) Number of Plasma cells produced, E) Affinity of plasma cells and F) Percentage of apoptotic cells. G-L) Dependence of GC volume kinetics and output, on cycling rates and antigen half-life for different antigen concentrations: AUC of the GC volume kinetics (G and J), Number of plasma cells (H and K) and Plasma cell affinity (I and L) observed on day 21 of the GC reaction. Legend shows average time spent by antigen particles in the interior (I) and surface (S) of FDCs (in minutes). Total antigen portion is 3000 and 1000 in each FDC for A-I and J-L, respectively). GC: Germinal Center; FDC: Follicular Dendritic Cell, PC: Plasma cell, AUC: Area Under Curve.

### Blocking antigen cycling can shut down GC reactions

We simulated the effect of blocking antigen cycling on GC reactions. When both, internalization and externalization of antigen were blocked at different time points of the GC reaction, shutdown of GC reaction was accelerated (Figure 4A) and plasma cell production was decreased (Figure 4C). Blockade of antigen cycling slightly impaired affinity maturation (Figure 4D). The impact on the GC reaction varied depending on the time point of blockade. Blocking internalization of antigen alone did not have any impact on the GC reactions (not shown). Blocking antigen externalization alone results in the accumulation of antigen in the interior of FDCs and results in a quick termination of GC reactions (supplementary Figure S2) when compared to blocking both internalization and externalization. These results suggest that antigen cycling can also be blocked to terminate GC reactions in a way similar to the effect of antibody feedback (46–48).

**Figure 4:**
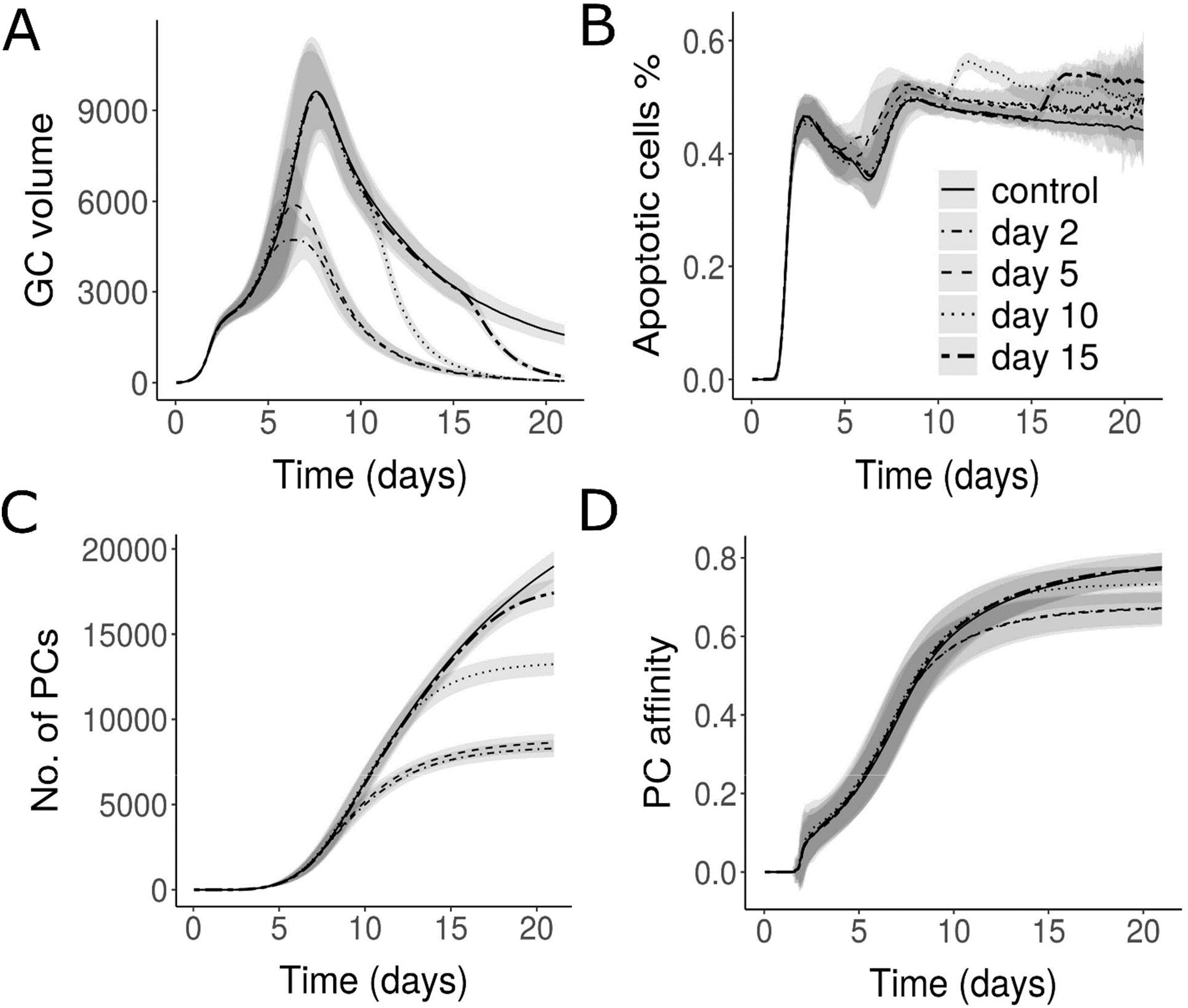
Effect of blocking both antigen internalization and externalization: A) GC volume, B) Percentage of apoptotic cells, C) Number of Plasma cells and D) Affinity of Plasma cells. Legend represents the time of blockade. GC: Germinal Center; PC: Plasma cell.

## Discussion

Discovery of IC-cycling in FDCs (13) has shed light on the previously unexplained observation of extended half-life of FDC antigen. In this study, we demonstrate the importance of IC-cycling in GC reactions by performing *in-silico* simulations. Our findings suggest that understanding the cycling of ICs in FDCs could have numerous implications in modulating GC reactions for efficient response towards vaccination, dissolving chronic GCs and for devising strategies to target pathogens such as HIV that remain protected in the FDCs (14).

Analysis of published experimental results (13), revealed the time scale of IC cycling in FDCs which was previously unknown and predicted that PE-IC particles spend approximately 21 minutes on the FDC surface and 36 minutes in the interior of FDCs. In the predicted range of antigen display kinetics, GC simulations showed robust dynamics and output production suggesting that cycling might not impact the GC dynamics directly, in the presence of high total antigen concentration. Consistently, changes in immune complex deposition in experiments did not lead to large alterations in the GC responses (49).

Considering that antigen integrity might be affected by prolonged exposure on FDC surface, our simulations demonstrate how antigen cycling could protect antigen degradation by minimizing surface exposure and at the same time allowing for efficient antigen uptake by GC B cells. Experimental data on time course of the number of GCs suggests that GCs are formed for at least until day 12 after immunization (50). Particularly, such late initialized GCs might be highly impaired if antigen is not retained stable for a long period.

According to our results, protection of antigen from degradation was largely dependent on the cycling rates and degradation is minimum if the antigen tends to spend longer time in the interior of FDCs. With limiting total antigen concentration, cycling rates determined the trade-off between protecting antigen from damage and allowing the efficient uptake of antigen by GC B cells. It is currently unknown whether ICs other than PE-ICs have a different cycling kinetics. However, antigen tagged to nanoparticles of different sizes differ in the distribution of these particles between the FDC surface and interior (15). Dynamics of transport of antigen to the lymphoid follicles (51–53), retention and presentation on FDCs vary with the nature and size of antigen particles (15) and these observations could be employed to engineer nanoparticles promoting efficient antigen retention in the FDCs in the context of vaccination (15). Consequently, studies on how antigen cycling varies for different antigen particles might also be useful in engineering antigens to enhance GC responses and improve the success of vaccination.

Moreover, blocking IC cycling prevents the efficient uptake and acquisition of antigen by FDCs (13). Hence, the importance of antigen cycling in FDCs on GC reactions could be more than what is estimated by our simulations. Further studies are needed to address the role of antigen cycling in initial acquisition of antigen by FDCs and its contribution to the efficiency of GC and memory responses. In the light of several studies showing that antigen delivery can be modulated to alter GC responses (21, 22), our findings contribute to understanding the dynamics of antigen within the FDCs brought about by cycling and its impact on the GC reaction kinetics. In addition to the GC reactions, long-term protection of antigen due to antigen cycling might be of importance in maintaining the memory responses. It has been shown that memory B cells reside close to contracted GCs (54) and it is thought that they might be reactivated upon exposure to antigen to maintain the serum antibody concentration levels (17, 55).

In the simulations, antigen cycling blockade at different time points resulted in termination of GC reactions. However, as the precise mechanism of antigen cycling is currently unknown, understanding the mechanism and gaining more insights into the factors regulating antigen cycling could help identify targets that might provide a therapeutic opportunity to dissolve antigen dependent chronic GCs. Munoz-Fernandez et al., showed that contractility of FDCs is enhanced by IL-2 and decreased by IL-10 suggesting cytokine-dependent changes in FDC morphology (56). IL-10 produced by Tfr (T follicular regulatory) cells (57) and a subset of human tonsillar follicular T cells (58) might be able to influence antigen display dynamics on FDCs in the GCs. Several mechanisms of action of Tfr cells in regulating GCs have been proposed including the direct inhibition of GC B cells and Tfh cells (59–61). In addition to these mechanisms, a potential impact of Tfr cells on FDCs and antigen display might also play a role. Moreover, a dynamic regulation of antigen cycling in the GCs during the course of the GC reaction could be expected depending on cytokine levels.

A number of factors including antibody feedback (46–48), CD40/CD40L interactions (62, 63) are able to influence GC shutdown even though the mechanism of shutdown is not well understood. As antigen was detectable in FDCs of lymphoid follicles even after the GC reaction ends (4), it is believed that the termination of GCs is not primarily due to the lack of antigen. However, future studies might explore whether natural alterations of antigen presentation by cytokines and other factors, could promote the termination of GC reactions.

## Contributions

TA, SB and MMH designed the study. TA performed simulations. SB and MMH supervised the project. TA, SB and MMH wrote the manuscript.

## Acknowledgements

We thank Michael Carroll and Balthasar Heesters for discussions and clarifying the experimental protocol. We also thank Sahamoddin Khailaie for constructive suggestions.

## Figures

**Figure S1:**
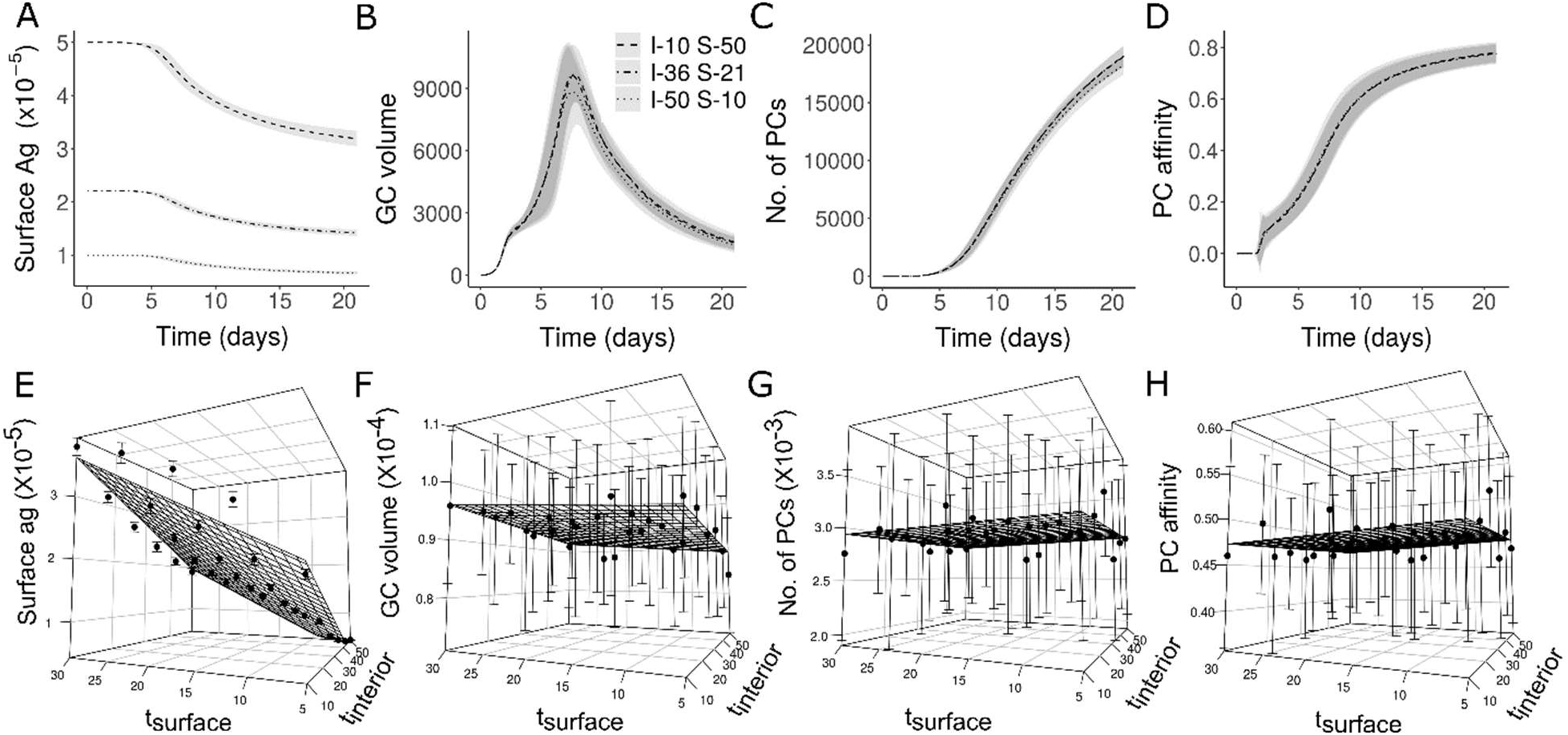
GC simulation as in Figure 2 with different cycling rates. (A-D) represents time course of GC readouts for 3 different cycling times and (E-H) GC readouts on day 8 for a range of cycling parameters: Surface antigen portions (A and E), GC volume calculated as number of GC B cells (B and F), Number of Plasma cells produced (C and G) and Affinity of plasma cells (D and H). Legend for A-D) shows average time spent by antigen particles in the interior (I) and surface (S) of FDCs (in minutes). Simulation readouts in (E-H) are shown as dots with error bars. The surfaces represent the regression planes calculated from the simulation data points. *t*_surface_ and *t*_interior_ are shown in minutes. GC: Germinal Centre; FDC: Follicular Dendritic Cell, PC: Plasma Cell. Initial antigen amount is 3000 antigen portions in each FDC.

**Figure S2:**
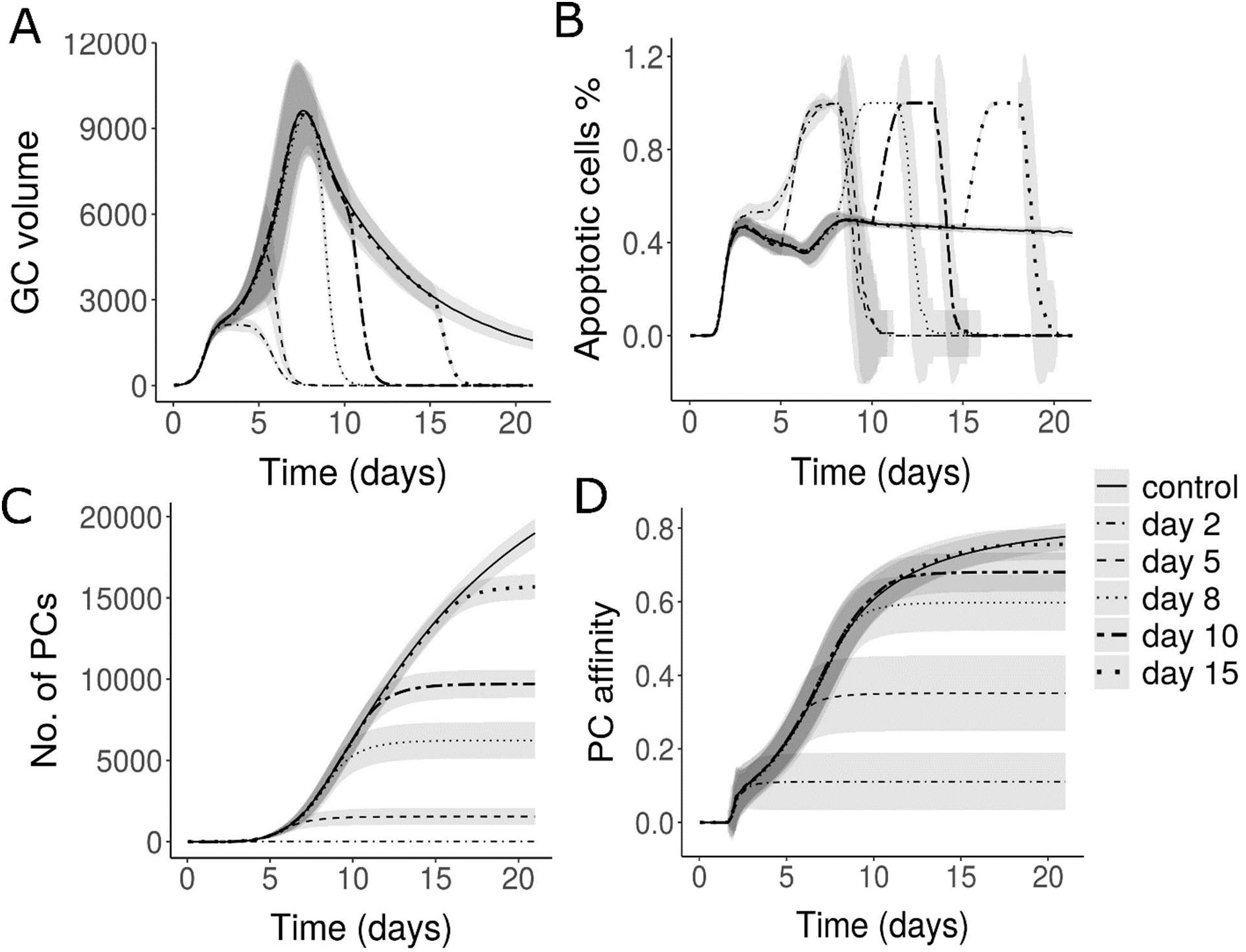
Simulations of antigen cycling blockade as in Figure 4 except that only the externalization of antigen is blocked: A) GC volume, B) Percentage of apoptotic cells, C) Number of Plasma cells and D) Affinity of Plasma cells. Legend shows the time of blockade.

